# Impact of evolutionary selection on dynamic behavior of MCAK protein

**DOI:** 10.1101/2020.12.04.412650

**Authors:** Shiwani Limbu

## Abstract

Kinesins of class 13 (kinesin-13s), also known as KinI family proteins, are non-motile microtubule binding kinesin proteins. Mitotic centromere-associated kinesin (MCAK), a member of KinI family protein, diffuses along the microtubule and plays a key role in microtubule depolymerization. Here we have demonstrated the role of evolutionary selection in MCAK protein coding region in regulating its dynamics associated with microtubule binding and stability. Our results indicate that evolutionary selection within MCAK motor domain at amino acid position 440 in carnivora and artiodactyla order results in significant change in the dynamics of *α – helix* and loop 11, indicating its likely impact on changing the microtubule binding and depolymerization process. Furthermore, evolutionary selections at amino acid position 600, 617 and 698 are likely to affect MCAK stability. A deeper understanding of evolutionary selections in MCAK can reveal the mechanism associated with change in microtubule dynamics within eutherian mammals.

## Introduction

Cell contains a cytoskeletal structure called microtubule which is important for maintaining its shape by providing structural support and also mediates the transportation of various molecules across the cell. Cytoskeleton is composed of highly dynamic polymer called microtubules made from the polymerization of tubulin dimers. The constant growth and shrinking of the microtubule ends help in maintaining various important cellular processes such as cargo transport, cytoskeleton adaptation, cell motility and cell division. The shrinking and elongation of microtubules are mediated by various proteins which are essential for cell migration and transportations. The KinI proteins belong to the microtubule depolymerases which helps in regulating the microtubule dynamics and depolymerization mechanism (Ems-McClung and Walczak, 2010). KinI family members are essential for managing cell polarity, dissociation of microtubule fiber during cell division, cell polarity and chromosomal segregation (Walczak et al., 2002). KinI proteins stabilizes the curved protofilament at the end of the microtubule polymer, thereby regulating the polymerization and disassembly process (Gardner et al., 2011). The Kin I proteins are an unusual when compared to other kinesin related proteins. KinI proteins do not move along the microtubule fiber in the presence of ATP, but instead bind to the end of microtubule to regulate its depolymerization activity (Desai et al., 1999).

The KinI family includes MCAK protein, which is essential for MT depolymerization activity. Inhibition of MCAK has shown to interfere with the dissociation and polar movement of chromosomes during the anaphase due to the loss of MT depolymerization (Maney et al., 1998). Furthermore, a loss of MTs in interphase as well as in mitotic cells indicates the role of MCAK in regulating MT depolymerization outside mitosis (Kinoshita et al., 2001; Maney et al., 2001, 1998). MCAK binds to the plus end of microtubule to initiate the depolymerization process. Various kinesin proteins undergo walking motion across the MT fiber, whereas MCAK diffuses on the microtubule end in ADP bonded state and initiates the depolymerization in ATP bonded state. Binding to the MT end initiates the dissociation of ADP molecule in exchange for ATP, allowing MCAK to bind tightly to the MT end (Belsham and Friel, 2017). ATP bound MCAK enhances bending of terminal tubulin dimer and thereby initiating the MT depolymerization process. A series of process involved in MT depolymerization as primarily regulated by series of phosphorylation reactions.

Structurally, MCAK contains an N-terminal domain (NT) which regulates its targeting to the centromere, a class-specific neck domain which is essential for MT depolymerization, a central catalytic motor domain which binds to the end of MT and regulates ATP hydrolysis, and a C-terminal domain (CT) which regulates the dimerization of MCAK, its catalytic activity, and spindle pole localization (Zong et al., 2016). The catalytic motor domain and the neck domain together for the minimal domain (MD) necessary to initiate the MT depolymerization activity in-vitro (Maney et al., 2001). The NT domain helps in localization of MCAK to the kinetochore where it binds to Sgo2 (Talapatra et al., 2015). NT domain also assists in MCAK localization to the plus end of MT allowing its binding to the end binding (EB) proteins (Talapatra et al., 2015). Moreover, the last 9 residues within the C-terminus of MCAK also play a significant role in its MT plus end localization (Moore et al., 2005). It has been shown that the removal of the last 9 amino acid residues at the C-terminus of MCAK leads to the auto-inhibition of its MT depolymerase activity (Moore and Wordeman, 2004). Residue segment from the end of motor domain to the end of CT domain (residues 584–725) has been reported to regulate the dimerization of MCAK as well as in its NT domain interaction (Maney et al., 2001). MCAK undergoes closed to open conformational transition upon its binding to the MT (Talapatra et al., 2015).

Evolution of MCAK protein has not been studied extensively so far. Moreover, there is a large gap of knowledge on evolution driven changes in MCAK and its impact on structure, function and dynamics. The role of MCAK in initialing microtubule depolymerization makes it more critical to understand the influence of evolution on the microtubule depolymerization mechanism. In this work we have estimated the positive and episodic diversifying selection to locate the amino acid changes occurred in MCAK through the coarse of evolution. Amino acid changes can affect the conformational and dynamic property of a protein molecule. Changes in conformation as well as its dynamic behavior affects the biological function of a protein. Moreover, the flexibility of amino acid residues within a protein affects its interaction propensity with its biological partner and affect its allosteric behavior. Therefore, molecular docking and molecular dynamics simulations were performed to study the evolution induced changes in the dynamic behavior of MCAK protein and investigate its impact on microtubule binding affinity. Our results show that evolutionary changes in MCAK affects its stability and dynamics which are associated with microtubule depolymerization process.

## Method

### Data collection

Gene exon sequences for eutherian mammals and marsupials were obtained from NCBI GenBank (Benson et al. 2005). All the available gene transcript sequences were obtained from GenBank. tBlastN (Altschul et al., 1990) was used to obtain non-human taxa protein coding sequences. Coding sequence alignment was performed using muscle (Edgar, 2004) alignment program. Structure of MCAK protein and Tubulin dimer (PDB ID: 5MIO) was collected from PDB database (Berman et al., 2000).

### Phylogenetic Tree Reconstruction

Sequence partition and maximum likelihood methods were implemented to reconstruct the phylogeny of mammals using MCAK, TUBA1A and TUBB3 gene coding sequences. Maximum likelihood (ML) phylogenetic analyses were performed using the GTRGAMMA method implemented in RAxML (Stamatakis, 2014). Partitionfinder (Lanfear et al., 2012) with the hcluster method was used to select the optimal partitioning scheme for the phylogenetic tree construction. ML searches were conducted using ten distinct randomized maximum parsimony starting trees. Phylogeny tree was visualized using FigTree (Rambaut, 2014).

### Test of pervasive positive selection

Site models allow ω to vary across sites but not across branches in the phylogeny. We implemented M1a and M2a site models to detect the evidence of positive selection in MCAK genes. Model M1a and M2a were implemented to test the evidence of pervasive positive selection in MCAK protein coding sequence. Model M1a allows protein coding sites to fall into purifying (ω <1) and neutral evolution (ω = 1), whereas M2a adds an additional category called positive selection with ω >1. (Nielsen and Yang, 1998). Here ω represents the ratio of nonsynoymous (dN) and synonymous (dS) substitution rates within the alignment. PAML (Yang, 2007) was used to generate ML estimates of parameters to detect sites evolving under positive selection.

HyPhy (Kosakovsky Pond et al., 2005) FEL (Fixed Efects Likelihood) (Kosakovsky Pond et al., 2008) model was implemented to estimate the rate of purifying selection in MCAK proteins. FEL uses a ML to estimate the dN and dS substitution rates for each alignment site using the corresponding phylogeny tree. Selection pressure at all the site across all the positions in the phylogeny tree is assumed to be constant in FEL model. dN and dS substitution rates are estimated by fitting MG94xREV model to each codon site after optimizing the tree branch lengths. Likelihood ratio hypothesis testing is then implemented to check if dN is significantly greater than dS.

SLAC (Kosakovsky Pond and Frost, 2005) (Single-Likelihood Ancestor Counting) implements a ML and counting appraoch to estimate the dN and dS substitution rates for each site of alignment. Branch lengths and the nucleotide substitution rates within the alignment and phylogeny tree are optimized under the MG94xREV model. ML is then used to find the ancestral sequence for all the within the phylogeny tree. dN and dS rates at each site are then counted using a modified version of the Suzuki-Gojobori counting method. Extended binomial distribution is used to estimate the significance of SLAC results.

### Test of episodic diversifying selection

Episodic diversifying selection test were performed using MEME (mixed effects models of evolution) (Murrell et al., 2012) implemented in HyPhy package. The MEME model allows variable ω values across all the lineage for a codon site under test, where each site is treated as a fixed component of model comparing ω ≤ 1 (proportion p) and ω+ (unrestriced, proportion 1-p). Here ω ≤ 1 indicates sites evolving neutrally or under the influence of negative selection, whereas ω+ indicates the evidence of diversifying selection for a given codon site. It computes the diversity within the phylogenetic branches differently as compared to the estimation of diversity among different branches in the phylogeny tree. Moreover, it estimates the measure of radiating selections that may indicate the association of selection events with the rapid change in the habitat.

### Molecular modelling and dynamics

N440S mutation was incorporated in the MCAK protein structure (PDB ID: 5MIO, chain C) using Pymol (DeLano, 2020). Molecular Dynamics Simulation (MDS) was conducted using Gromacs (Hess et al., 2008; Pronk et al., 2013) to study the impact of evolution on MCAK protein dynamic behavior. Structure of native and N440S mutant MCAK was used as starting structure for MDS. Energy minimization was conducted using conjugate gradient method. Temperature within the box was controlled using Berendsen temperature coupling method (Berendsen et al., 1984). Particle Mesh Ewald method (Cheatham et al., 1995) was implemented to compute the electrostatic interactions within the protein. 5 nanoseconds (ns) of position restrained simulation and 700 ns of unrestrained MDS was conducted to obtain the motion trajectory of native and the mutant structure. RMSD and RMSF analysis were conducted to estimate the change in fluctuation among the native and the mutant structure. Total energy trajectory was estimated to study the change in stability of protein. Number of distinct hydrogen bonds (NHbonds) was calculated to analyze the molecular changes within the protein structure. All graphs were plotted using Grace GUI toolkit.

### SNP analysis

I-Mutant 2.0 (Capriotti et al., 2005) was implemented to test the impact of evolution driven mutational changes on MCAK stability. It was trained on mutation induced free energy change values taken from ProTherm database (Bava et al., 2004). I-Mutant 2.0 reports mutation induced likely changes in free energy values of the protein.

### Molecular docking

Molecular docking of MCAK with tubulin dimer was performed using HADDOCK tool (De Vries et al., 2010; Dominguez et al., 2003). Total interaction energy, buried surface area, van der Waals energy, restraints violation energy, electrostatic energy and desolvation energy were used to estimate the impact of change in MCAK tubulin dimer binding affinity change induced by evolutionary changes in amino acid residues.

## Results and Discussion

The non-synonymous to synonymous rate of substation ratio (ω) can be implemented to test selection on the protein coding gene sequence. ω > 1 suggests positive selection pressure leading to the fixation of an amino acid change through the coarse of evolution whereas ω < 1 indicates the presence of purifying selection in the coding region of a gene. Moreover, ω ~ 1 indicates the presence of neutral selection at the given position on gene. Purifying selection represents a tendency of a gene to preserve the existing amino acid from any non-synonymous changes. Errors in sequencing and annotation can influence the codon position and subsequently effect the amino acid sequence and are likely to affect synonymous as well as the nonsynonymous sites. Therefore, the quality of sequence alignment data can not only affect the reconstruction of phylogeny tree, but it can also lead to the overestimation of the ratio of dN and dS leading to the overestimation of pervasive positive selection. We implemented sequence annotations checks and sequence alignment refinements to improve the quality of sequence data used in this study. We found that the phylogeny tree constructed in this work was nearly congruent to the topology of mammalian phylogeny tree published in Meredith et al. 2011 (Meredith et al., 2011).

Evolution driven selection in residue position can influence the functional and dynamic behavior of the protein. Using our phylogenetic tree [Fig. 1], we tested for positive selection in the protein coding region of MCAK. Given the intriguing pattern of positive selection observed in MCAK protein coding region [Table 1-3], it is likely that evolution may have played a significant role in shaping the function and dynamic behavior of protein. 11 amino acid positions were found to be positively selected in the estimation performed using PAML [Table 1]. Out of these 11 positively selected positions, 5 residue position (145, 440, 600, 617 and 698) were commonly found to be positively selected in SLAC [Table 2] and FEL [Table 3] results. Positive selection at 600, 617 and 698 may change the behavior of C-terminal mediated MCAK regulation. As discussed earlier, CT to the motor domain (residues 584–725) helps in MCAK dimerization and NT domain interaction independently of the motor region. Moreover, nine amino acid residues within the CT domain (_709_QLEEQASRQISS_720_) of MCAK is shown to play an important role in plus tip tracking. We observe change from Glutamine (polar uncharged amino acid) to Arginine (Aromatic positively charged amino acid residue) at residue position 698. Such large change in amino acid property closed to CT domain may influence the MCAK dimerization and microtubule binding activity. We observed a slight loss of stability, with −0.68 free energy change, induced by R698N, Furthermore, non-polar to polar residue change at I600T and Proline (restricted side chain rotations) to Threonine at P617T may influence MCAK C-terminal residue dynamics. I600T and P617T induces a large change in free energy value (−1.65 and −1.70 respectively) in MCAK protein. Such large change may influence the dynamics of C-terminal region at a larger extend. V145I may not influence the dynamics of MCAK C-terminal domain due to the similarity in biophysical property and side chain length of Valine and Isoleucine residues. V145I induces negligeable change in free energy value (−0.02), indicating a negligeable change in stability of protein.

**Fig 1.**
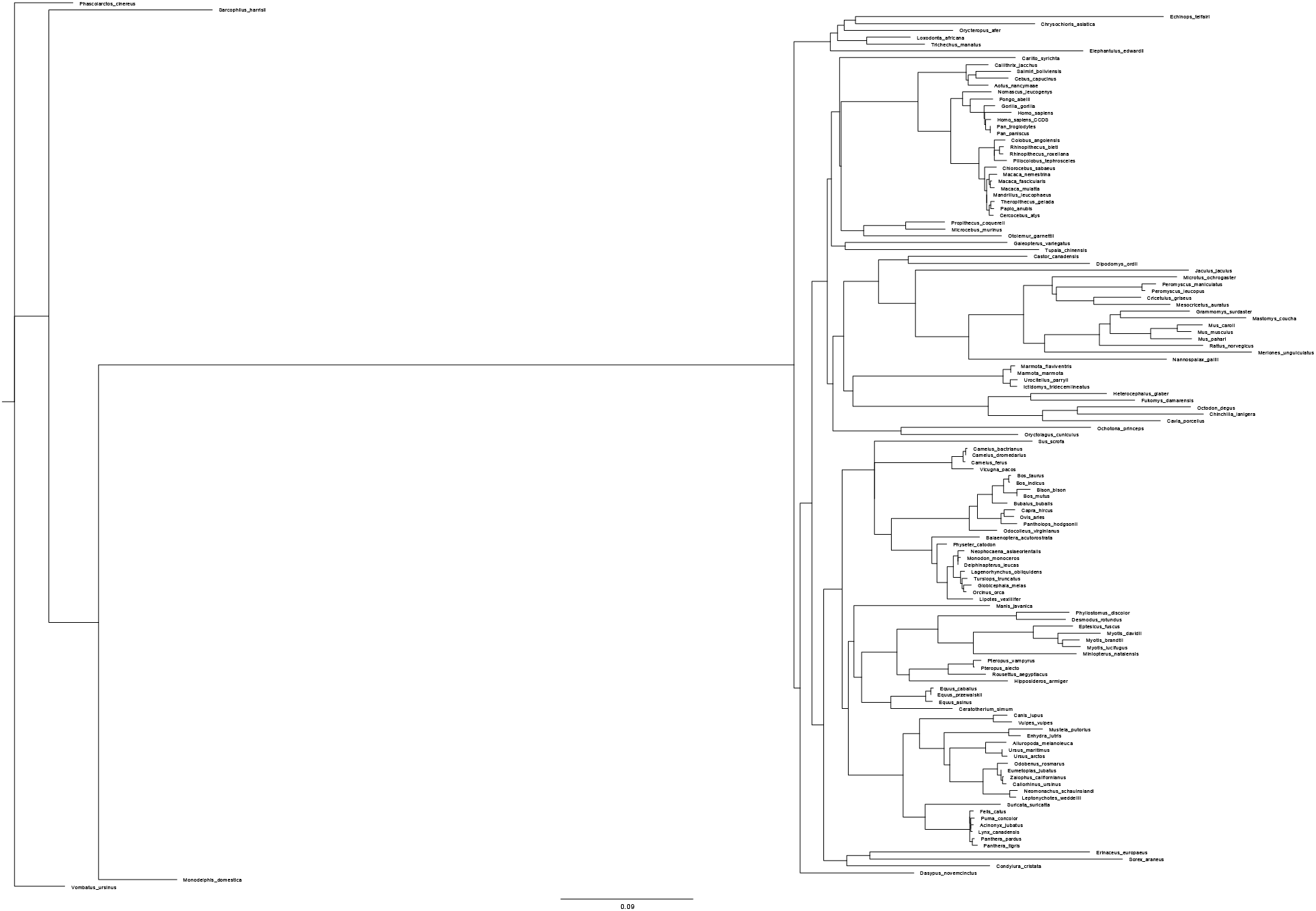
Mammalian phylogeny tree constructed using coding sequence data of MCAK, TUBA1A and TUBB3 protein.

**Table 1.** Pervasive positive selection in MCAK protein estimated using PAML.

| Amino acid site | Pr(w>1) |
| --- | --- |
| 2 | 0.596 |
| 118 | 0.762 |
| 120 | 0.721 |
| 124 | 0.649 |
| 136 | 0.686 |
| 145 | 0.871 |
| 150 | 0.671 |
| 440 | 0.902 |
| 600 | 0.971 |
| 617 | 0.952 |
| 698 | 0.954 |

**Table 2.** Pervasive positive selection in MCAK estimated using SLAC.

| Amino acid site | ES | EN | dN-dS | P-value |
| --- | --- | --- | --- | --- |
| 145 | 1.06 | 2.14 | 1.89 | 0.0376 |
| 440 | 0.725 | 2.30 | 1.61 | 0.0284 |
| 600 | 0.875 | 2.21 | 2.18 | 0.0448 |
| 617 | 1.03 | 2.04 | 2.52 | 0.0211 |
| 698 | 0.894 | 1.46 | 3.35 | 0.00143 |

**Table 3.**
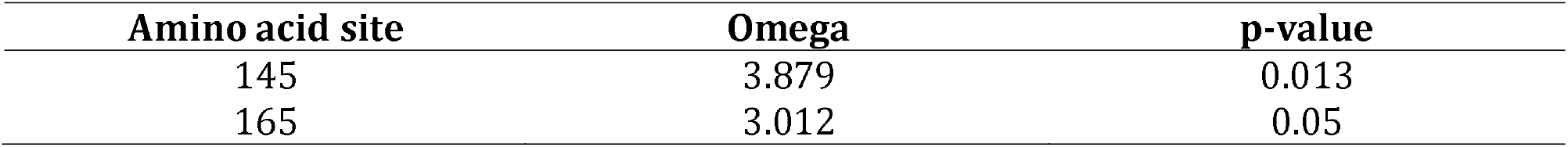
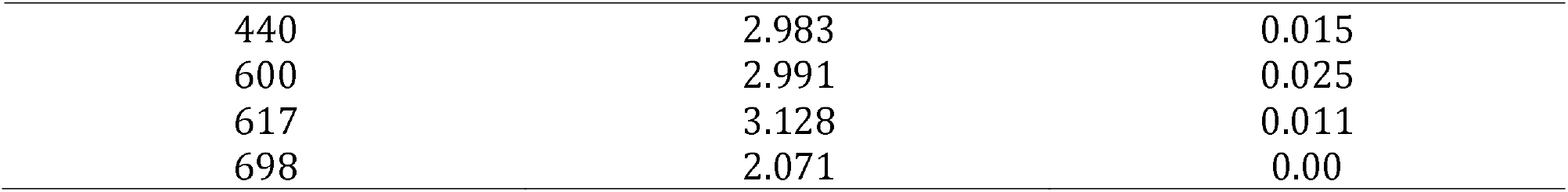
Pervasive positive selection in MCAK estimated using FEL.

Our results indicated an evidence of positive selection [Table 1-3] as well as divesifying selection [Table 4] at residue position 440 in the motor domain of MCAK protein. Majority of species within order carnivora and artiodactyla contains serine amino acid residue at position 440, whereas primata, rodentia as well as chiroptera consists arginine amino acid residue at position 440. We also observed arginine amino acid residue within marsupials animals, indicating the recent occurrence of N440S selection event in eutherian carnivora and artiodactyla order. Presence of a selection event in MCAK motor domain (N440S) was investigated for its effect on dynamics using molecular dynamics simulation. Selection even N440S will be referred as mutant in this manuscript. To observe the effect of selection N to S on the dynamic behavior of residues, the RMSF values of native and the mutant (N440S) MCAK protein structure were computed from the MDS trajectory (Fig. 2). Results indicated higher degree of flexibility in the mutant as compared to the native MCAK structure. Our results indicate that the selection changes the dynamics of MCAK Loop-11 and *α – helix*. Loop-11 and *α – helix* are important for microtubule end recognition and helps MCAK to distinguish between the lattice of microtubule from its end. Both Loop-11 and *α – helix* plays an important role in microtubule depolymerization (Patel et al., 2016). The microtubule end assists in activation of the MCAK ATPase activity by the dissociation rate of ADP molecule. Mutation in *α – helix* has been shown to reduce the ability of MCAK to stay at the end of microtubule for a longer duration as well as in affecting the ability of microtubule end to accelerate dissociation rate of ADP molecule from the MCAK motor domain [38]. N440S induced change in dynamics of MCAK *α – helix* and residues in close proximity to Loop-2 indicates adaptive changes in the activity of MCAK motor domain. An adaptive selection in the MCAK *α – helix* may have helped in evolution of microtubule depolymerization rate in Eutherian animals. A severe change in dynamics of 10 residues (267-277) were observed in the RMSF plot. Segment 267-277 is in close proximity to the ATP binding pocket and may influence the hydrolysis of ATP molecule and ADP dissociation, thereby affecting the microtubule depolymerization rate.

**Fig 2.**
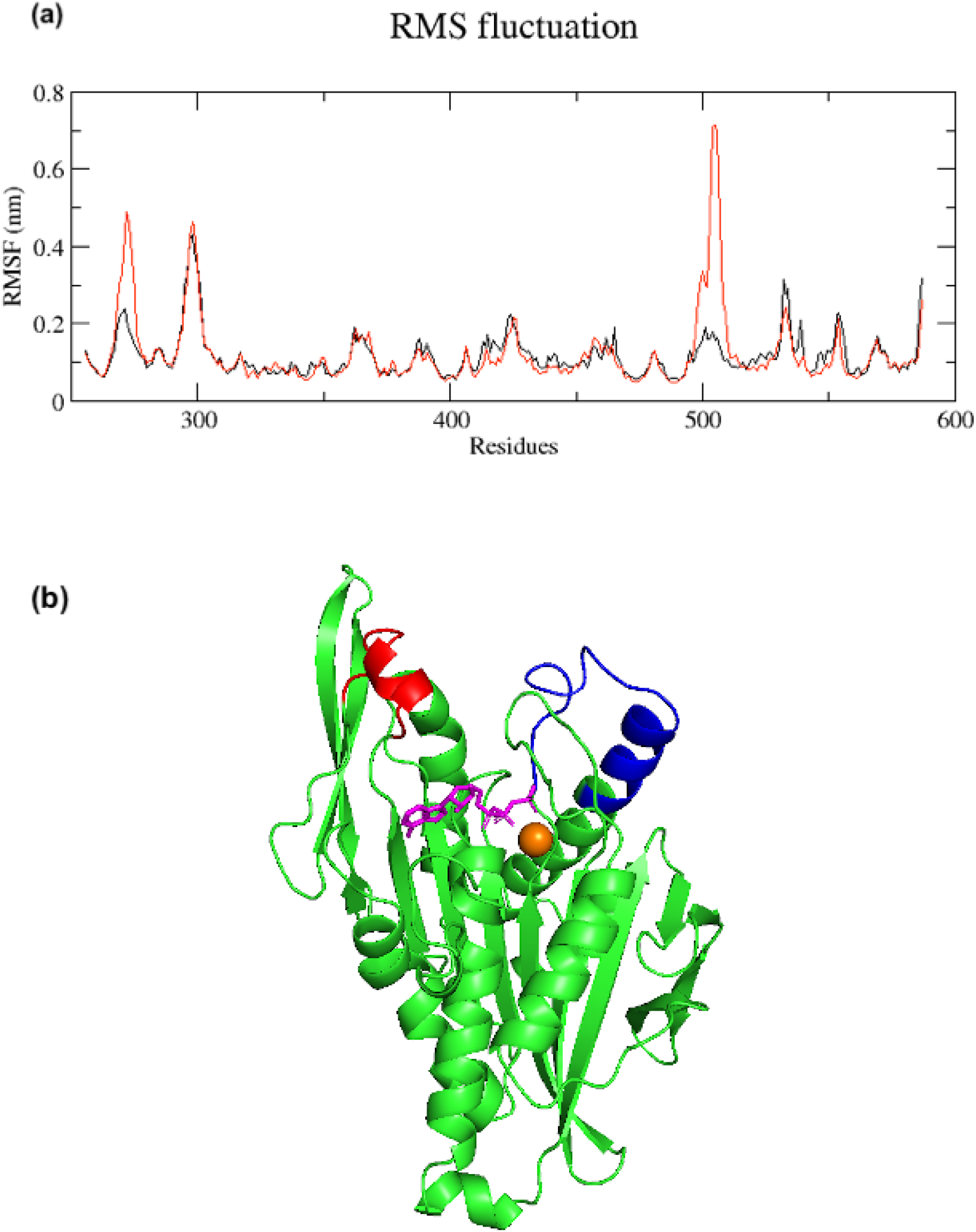
a) RMSF plot of native (black) and mutant (red) MCAK protein. The peak changes in RMSF are observed in residue position 267-277, which is a part of Loop-11 and position 494-514 which is a part of of MCAK protein, b) residue positions 267-277 is shown in red and residue position 494-514 is shown in blue.

**Table 4.** Diversifying selection event in MCAK estimated using MEME.

| Protein | Amino acid site | $\alpha$ | $\beta^-$ | $p^-$ | $\beta^+$ | $P^+$ | LRT | p-value |
| --- | --- | --- | --- | --- | --- | --- | --- | --- |
| MCAK | 28 | 0.77 | 0.14 | 0.98 | 87.96 | 0.02 | 12.27 | 0.00 |
|  | 88 | 1.29 | 0.45 | 0.99 | 346.62 | 0.01 | 6.85 | 0.01 |
|  | 150 | 0.71 | 0.00 | 0.79 | 6.97 | 0.21 | 8.97 | 0.00 |
|  | 152 | 1.31 | 0.00 | 0.90 | 8.34 | 0.10 | 5.03 | 0.04 |
|  | 180 | 0.88 | 0.19 | 0.99 | 125.65 | 0.01 | 9.73 | 0.00 |
|  | 440 | 0.00 | 0.00 | 0.11 | 0.68 | 0.89 | 5.94 | 0.02 |
|  | 479 | 0.24 | 0.07 | 0.98 | 29.46 | 0.02 | 9.90 | 0.00 |
|  | 582 | 0.82 | 0.00 | 0.99 | 28.57 | 0.01 | 5.32 | 0.03 |
|  | 583 | 0.00 | 0.00 | 0.97 | 2.31 | 0.03 | 5.82 | 0.02 |
|  | 600 | 0.51 | 0.51 | 0.27 | 1.96 | 0.73 | 5.04 | 0.04 |
|  | 620 | 0.92 | 0.52 | 0.98 | 313.33 | 0.02 | 11.50 | 0.00 |
|  | 621 | 0.11 | 0.00 | 0.92 | 11.10 | 0.08 | 14.97 | 0.00 |
|  | 663 | 0.83 | 0.00 | 0.99 | 55.70 | 0.01 | 9.61 | 0.00 |
|  | 700 | 0.22 | 0.00 | 0.94 | 6.25 | 0.06 | 8.75 | 0.01 |
| TUBA1A | 234 | 0.54 | 0.00 | 0.99 | 513.50 | 0.01 | 6.10 | 0.02 |
|  | 357 | 0.00 | 0.00 | 0.99 | 3085.47 | 0.01 | 7.07 | 0.01 |
| TUBB3 | 7 | 0.27 | 0.00 | 0.99 | 9155.40 | 0.01 | 7.92 | 0.01 |
|  | 121 | 2.27 | 0.00 | 0.99 | 10000.00 | 0.01 | 5.67 | 0.00 |
|  | 189 | 0.00 | 0.00 | 0.98 | 29.67 | 0.02 | 12.56 | 0.00 |
|  | 275 | 0.16 | 0.00 | 0.98 | 38.58 | 0.02 | 16.24 | 0.00 |
|  | 332 | 0.33 | 0.00 | 0.98 | 67.59 | 0.02 | 18.74 | 0.00 |
|  | 346 | 0.34 | 0.00 | 0.99 | 62.93 | 0.01 | 10.87 | 0.00 |
|  | 351 | 0.55 | 0.00 | 0.96 | 18.97 | 0.04 | 12.98 | 0.00 |

Hydrogen bond plays a key role in maintaining the conformational stability and dynamics of a protein molecule. Changes in the number of hydrogen bond among native and the mutant structure was investigated from the MDS trajectory to study the impact of mutation on protein stability and associated conformational changes. Relatively lower number of hydrogen bonds were observed in the mutant when compared to the native (Fig. 3). Change in hydrogen bond formation can explain the relative change in RMSF values in *α – helix* of MCAK. Hydrogen bonds plays an important role in maintaining the thermostability of a protein structure. Hydrogen bond changes can affect the stability and therefore change the dynamic behavior of a protein molecule. To examine the impact of loss of hydrogen bonds in mutant protein, we estimated the change in energy spectrum throughout the simulation. Mutant indicated rise in total energy value when compared to the native MCAK structure, indicating gain of stability of native as compared to the mutant (Fig. 4). The changes in hydrogen bond and energy spectrum explains the dynamic changes observed in RMSF results in MCAK. Such changes may affect the binding of MCAK to microtubule and affect microtubule depolymerization rates.

**Fig 3.**
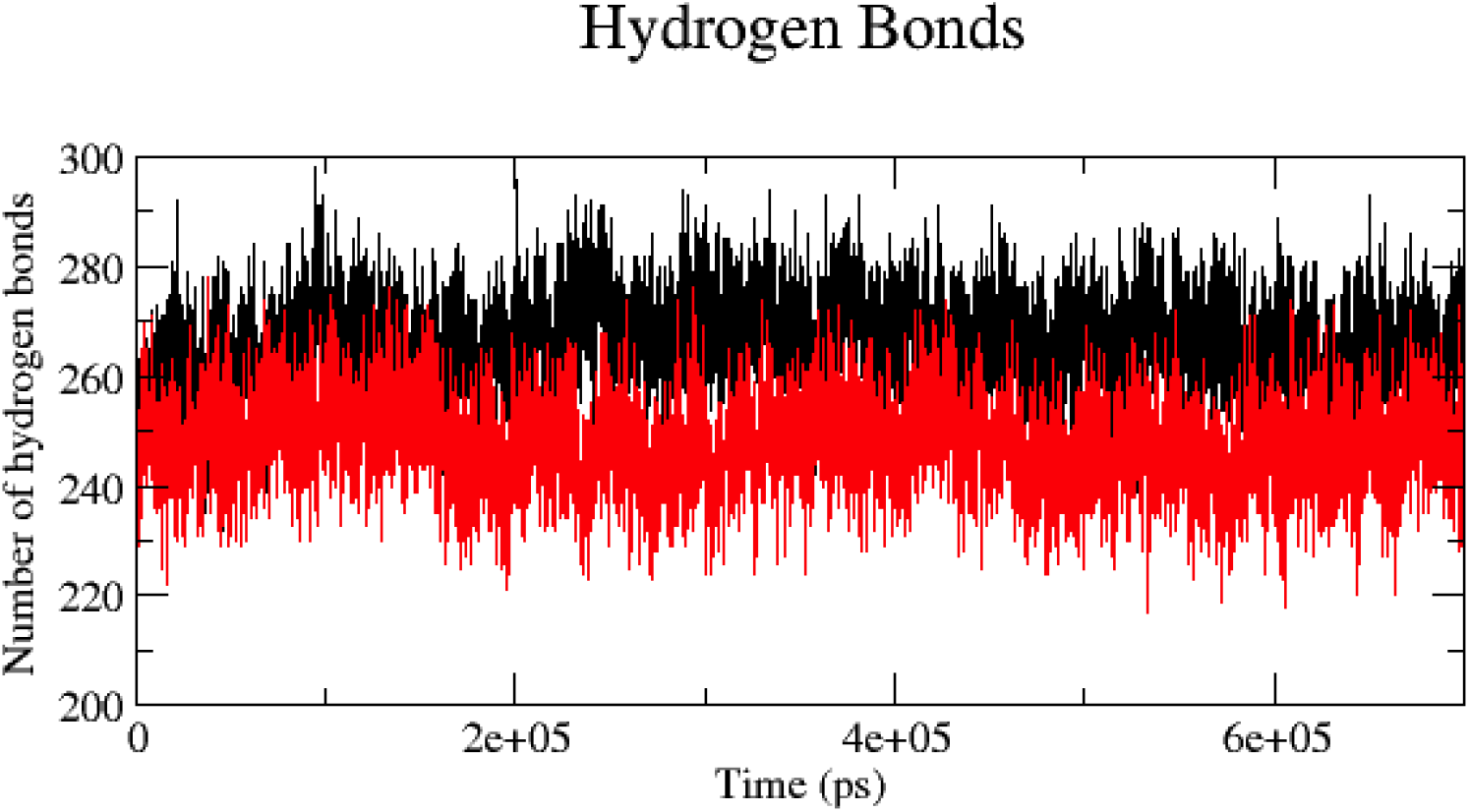
Hydrogen bond plot of native (black) and mutant (red) MCAK protein.

**Fig 4.**
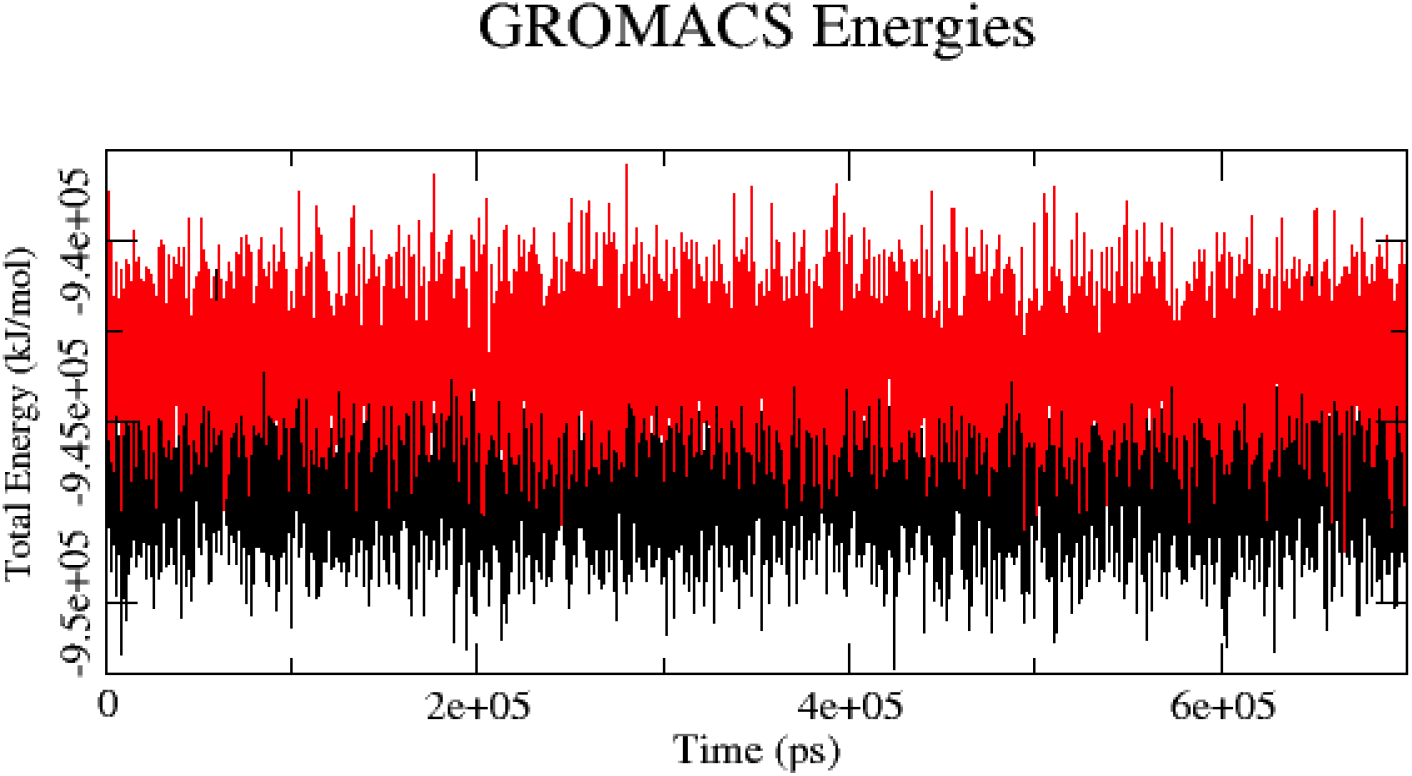
Total energy plot of native (black) and mutant (red) MCAK protein.

Molecular docking of tubulin dimer and MCAK was performed to estimate the impact of N440S selection event on their binding affinity to microtubules. Analysis of the native and mutant docked molecule revealed a notable difference between the binding ability of native and mutant MCAK protein with the tubulin dimer. van der Waals and electrostatic interaction energy played a key role in regulating the overall interaction energy between MCAK (native and mutant) and the tubulin dimer. Native MCAK-tubulin dimer bonded structure showed a van der Waals and electrostatic energy of −126.9 +/− 11.3 kcal/mol and −1143.3 +/− 40.9 kcal/mol respectively. Native showed a total native MCAK-tubulin dimer interaction energy of −286.0 +/− 8.4 kcal/mol. Mutant complex showed relatively high van der Waals and electrostatic energies of −119.3 +/− 5.4 kcal/mol and −1083.5 +/− 49.5 kcal/mol, respectively, as well as relatively higher amount of total interaction energy −264.0 +/− 7.1 kcal/mol when compared to the native. Change in interaction energy values shows that evolution has mediated a change in MCAK binding affinity to the microtubule.

The measure of buried surface area (BSA) provides an estimate of intrinsic flexibility within a protein structure. Native MCAK-tubulin dimer complex showed BSA of 4498.9 +/− 96.7 whereas in mutant it was 4215.5 +/− 222.8. Native MCAK motor domain showed −41.24 Kcal/mol desolvation energy whereas mutant showed −30.35 Kcal/mol desolvation energy value. The values of desolvation energy and BSA were in concordance to the docking energy values of native and mutant. Significant change in the interaction energy, BSA and desolvation energy were observed in mutant when compared to native, further indicating the role of N440S selection event in changing binding affinity to the microtubule within carnivora and artiodactyla.

## Conclusion

Role of molecular evolution in shaping the dynamic behavior of MCAK has not been studied before. Our results show that MCAK accumulated set of mutations in N-terminal, C-Terminal as well as in the motor domain. Dynamic analysis indicated that the evolution in motor domain may have changed the binding of MCAK to microtubule and it has also changed the dynamic behavior of MCAK in *α – helix* and loop 11, indicating its impact on microtubule depolymerization. Amino acid residue changes in C-terminal domain showed a high extend of change in stability of the protein. Our results indicate that evolution has helped in changing the dynamic behavior of MCAK protein in the segments which plays a key role in initiating the depolymerization of microtubules. We further aim to investigate the impact of these evolution driven changes MCAK at a cellular level.

## Acknowledgement

I thank Dr. Rituraj Purohit, Senior scientist, Council of Scientific and Industrial Research for his valuable suggestions in this work.

## Abbreviations

MT: Microtubule
MCAK: Mitotic centromere-associated kinesin
NT: N-terminal domain
CT: C-terminal domain
CCDS: Consensus CDS
FEL: Fixed Effects Likelihood
ML: Maximum Likelihood
SLAC: Single-Likelihood Ancestor Counting
dS: synonymous
dN: nonsynonymous
RI: reliability index
MEME: Mixed Effects Models of Evolution
NHbonds: Number of distinct hydrogen bonds

